# Dengue and Zika virus capsid proteins bind to membranes and self-assemble into liquid droplets with nucleic acids

**DOI:** 10.1101/2021.06.07.447421

**Authors:** Ernesto E. Ambroggio, Guadalupe Soledad Costa Navarro, Luis Benito Pérez Socas, Luis A. Bagatolli, Andrea V. Gamarnik

## Abstract

Liquid-liquid phase separation is prone to occur when positively charged proteins interact with nucleic acids. Here, we studied biophysical properties of Dengue (DENV) and Zika (ZIKV) virus capsid proteins to understand the process of RNA genome encapsidation. In this route, the capsid proteins efficiently recruit the viral RNA at the ER membrane to yield nascent viral particles. However, little is known either about the molecular mechanisms by which multiple copies of capsid proteins assemble into nucleocapsids or how the nucleocapsid is recruited and wrapped by the ER membrane during particle morphogenesis. Here, we measured relevant interactions concerning the viral process using purified DENV and ZIKV capsids proteins, membranes mimicking the ER lipid composition and nucleic acids at *in vitro* conditions. We found that both ZIKV and DENV capsid proteins bound to liposomes at liquid-disordered phase regions and docked exogenous membranes and RNA molecules. When the proteins bound nucleic acids, droplet liquid-liquid phase separation was observed. We characterized these liquid condensates by measuring nucleic acid partition constant and the extent of water dipolar relaxation observing a cooperative process for the formation of the new phase that involves a distinct water organization. Our data supports a new model in which capsid-RNA complexes directly bind the ER membrane, seeding the process of RNA recruitment for viral particle assembly. These results contribute to understand the viral nucleocapsid formation as a stable liquid-liquid phase transition, which could be relevant for Dengue and Zika gemmation, opening new avenues for antiviral intervention.

## Introduction

Proteins in solution can phase separate into liquid compartments by intermolecular interactions with nucleic acids. Such transitions occur with a first seed formed either in bulk or at the interface of substrates/soluble components, leading to a new liquid entity that display different physical properties compared to those of its surroundings. The nucleolus is the paradigm of these liquid compartments, nowadays known as membrane-less organelles, but already described in the 1830s [see 1, 2 and references therein]. It is accepted that one of the in-vivo/in-vitro interaction that triggers liquid-liquid phase separation is the association of RNA with proteins. One mechanism is by enabling multimerization of the RNA binding protein (RBP) (3). Furthermore, there are several reports indicating that proteins can also self-phase separate into liquid phases at the lipid bilayer interface (4). The capsid proteins (C) of Dengue (DENV) and Zika (ZIKV) virus interact with viral RNA, forming a nucleocapsid (NC), in the process of viral genome packaging. This process is called encapsidation and up to date a clear notion concerning the molecular mechanisms behind this phenomenon are not fully understood (5, 6).

DENV and ZIKV are relevant mosquito borne human pathogens of the Flavivirus genus. Upon infection, the positive stranded RNA genome is directly used as messenger for translating a large polyprotein, which is proteolytically processed at the ER membrane. The viral capsid is the first protein encoded in the open reading frame and remains inserted into the ER membrane by a transmembrane anchor peptide that links capsid to the pre-Membrane protein (prM). Two successive cleavages exerted by cellular and viral proteases are necessary for capsid release. The viral protease NS2B/NS3 is responsible of cleaving the anchor peptide, permitting the release of the mature capsid protein at the cytoplasmic side (5, 6). Capsid proteins recruit the newly synthesized viral genome to form the NC complex, which subsequently buds into the ER lumen gaining the lipid bilayer together with the viral proteins E and prM (5). How NC is recruited to the ER membrane is under extensive scrutiny. So far, high resolution microscopy techniques such as cryoEM do not provide compelling evidence that the C or NC are physically associated to the inner leaflet of the viral lipid membrane (7–9). To attempt to answer these questions, we use fluorescence methods to measure the interaction of both DENV and ZIKV capsids with ER-mimicking lipid membranes in the absence or in the presence of nucleic acids. We describe how ZIKV and DENV capsid proteins interact with liposomes and RNA molecules. We measured the protein-membrane binding and the ability of DENV and ZIKV capsids to recruit ssDNA or RNAs and liposomes onto the interface of giant unilamellar vesicles (GUVs). The capsid-RNA-membrane association was found at regions corresponding to liquid disordered (L_d_) phase. Also, the viral protein-RNA interaction generated phase-separated liquid droplets, with clear-cut changes on water organization within the droplets respect to their surroundings. We further characterized this process by computing an apparent dissociation constant for the nucleic acids/protein interaction, together with the determination of the cooperativity of the process.

## Results

### DENV and ZIKV capsid proteins bind and dock liposomes

To investigate whether DENV and ZIKV capsid proteins interact with membranes, we used different fluorescence methods. We first monitor the characteristics of the fluorescence emission spectra of a single tryptophan (W) contained in the proteins (highlighted in Figure 1A and D) in absence and presence of ERmix LUVs. W florescence emission properties are well known to depend on the polarity of the environment (10). The W emission spectra of both DENV and ZIKV capsids show a shift towards shorter wavelengths (Figure 1B and E) in the presence of LUVs, which is quantified by the ratio of the fluorescence intensities at 337 nm and 346 nm (Figure 1C and F). This indicates that upon membrane interaction the W senses a less polar milieu. The polarity change can be associated to a protein structural rearrangement, to a new environment due to the proximity of the amino acid to the membrane, or both. To answer this question, we took advantage of the well-known energy transfer effect between W and the NBD dye, represented in Figure 1G (11). The fluorescence intensity of NBD rises upon titration of NBD-dyed ERmix LUVs (where NBD is at the interface of the membrane) with increasing amounts of DENV and ZIKV capsid proteins, Figure 1H and I respectively. These data indicate that proteins bind to membranes and their W are sufficiently close to the lipid bilayer interface (2.2 nm; (12)) to transfer fluorescence energy to NBD.

**Figure 1.**
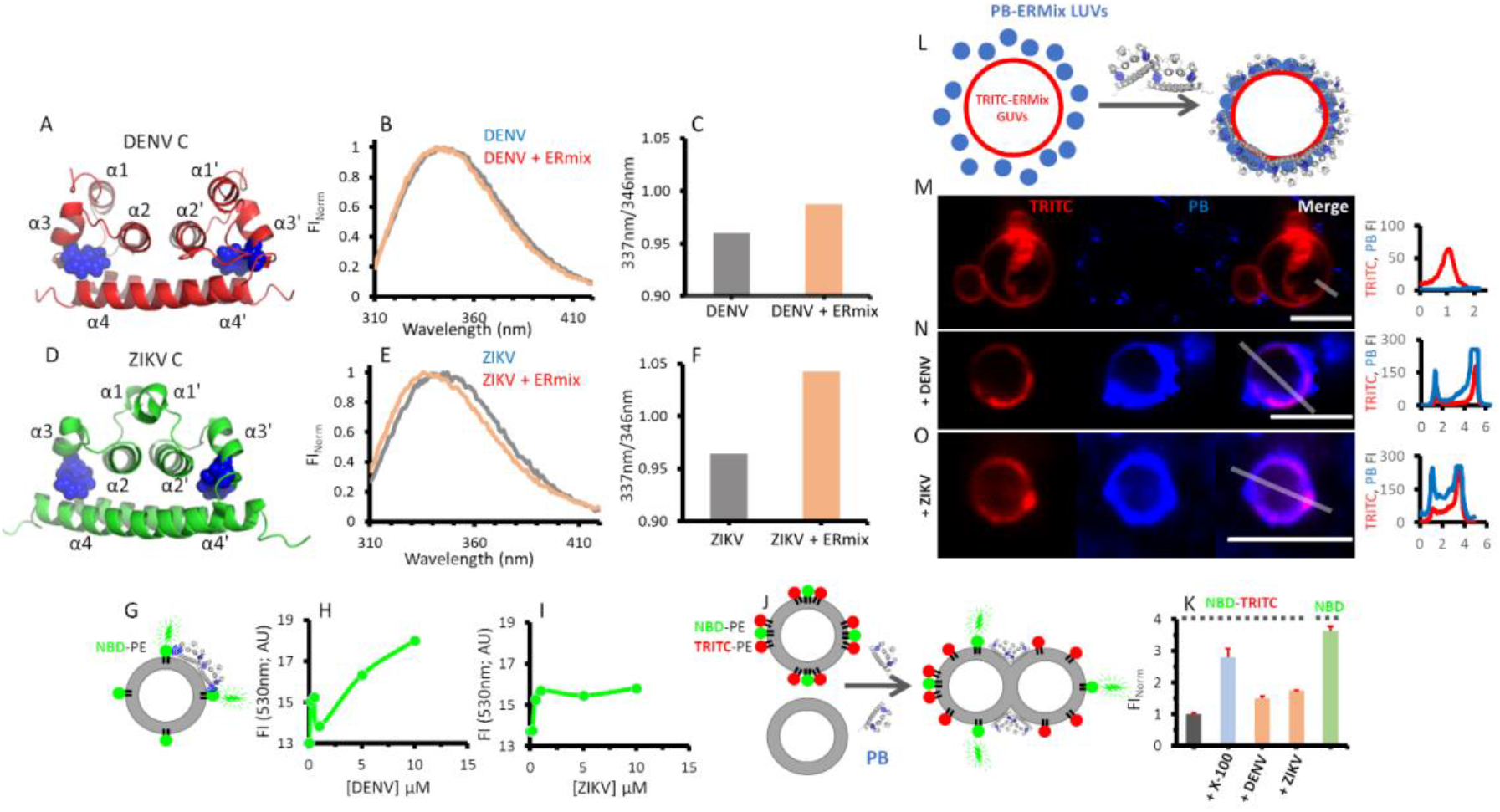
DENV and ZIKV capsid interaction with ERmix liposomes. DENV (A) and ZIKV (D) capsids dimeric structures where the W residue is represented in blue spheres. Tryptophan fluorescence emission spectra and quantification of the 337 nm/346 nm wavelength ratio for DENV (B, C) and ZIKV (E, F), respectively, in the absence (grey) and in the presence (light brown) of ERmix LUVs. Cartoon representing W-NBD FRET (G) when capsid proteins interact with NBD-PE labelled ERmix LUVs. The increase of NBD fluorescence intensity (FI) due to W proximity (λ_ex_ = 280 nm; λ_em_ = 530 nm) is measured in function of DENV (H) or ZIKV (I) addition. The lipid mixing assay represented in J where NBD fluorescence (donor dye) increases when lipid content from nonlabelled LUVs fusses with NBD-PE/TRITC-PE dyed LUVs. The normalized fluorescence intensity of NBD of the mixture NBD-PE/TRITC-PE stained ERmix LUVs + ERmix LUVs alone for the different conditions is showed in (K). A GUVs-LUVs docking assay triggered by protein addition is schematized in L where blue-fluorescent LUVs are recruited to the membrane of red fluorescent GUVs after addition of the capsid proteins. Confocal fluorescence images of the ERmix GUVs labelled with TRITC-PE (red channel; TRITC) in the presence of ERmix LUVs doped with PB-PE (blue channel, PB) in the absence (M) or in the presence of DENV (N) or ZIKV (O) capsids are shown. From the color merge image (Merge) the TRITC (red line) and PB (blue line) fluorescence intensity profile was measured along the drawn grey line and plotted at the right of each micrography. Note that the PB fluorescence signal profiles plotted in N and O reach their maximum values since image are acquired with exactly same settings as the control (M). Results are from at least two independent experiments (quantitative plots showing average and standard deviations) and images show representative observations. Scale bar in confocal images is 5 µm.

During the above-mentioned experiments, a change in the turbidity of the sample was observed, suggesting that protein interaction with the liposomes may induce a proteo-liposome aggregation. To further explore whether this process involved lipid mixing between different liposomes (i.e., throughout fusion or hemifusion of LUVs), we performed a FRET assay using the NBD-PE/TRITC-PE couple, as schematized in Figure 1J (see material and methods for more details). Upon addition of either DENV or ZIKV capsid proteins to the labelled/non-labelled liposome mixture, an increase of NBD fluorescence signal about 1.5 times higher than the control (normalized to 1) was observed, indicating lipid exchange among LUVs. Interestingly, this change represents on average half of the full intensity registered respect to controls (Figure 1K). These data support the idea that hemifusion is occurring upon protein-membrane interaction. In order to achieve spatially resolved information about this interaction, we used confocal fluorescence microscopy to visualize directly the membrane docking between ERmix giant unilamellar vesicles (GUVs) and ERmix LUVs. For this, TRITC-labelled GUVs were incubated with pacific-blue stained LUVs in the absence or presence of viral capsid proteins. In the absence of proteins, there was no LUVs signal at the membrane of GUVs (Figure 1M). However, upon addition of DENV (Figure 1N) or ZIKV (Figure 1O) capsids, there is a clear recruitment and enrichment of LUVs at the GUVs membrane interface, where a strong spatial co-localization of the two dyes was observed.

### DENV and ZIKV recruit liposomes and nucleic acids at the GUV membrane interface

DENV and ZIKV capsids form the NC by binding to the viral RNA genome, however, it is still unknown how the NC is recruited into the nascent viral particle at the ER membrane. The NC must get in close contact with ER membranes for particle morphogenesis, but it is uncertain whether capsid-lipid membrane interaction is a driving force in this crucial step. To further analyze the role of the interaction with lipid membranes of the DENV and ZIKV capsid proteins, we carried out an experiment where ERmix GUVs were incubated with a fluorescently labelled ssDNA or ssRNA in the presence or absence of such proteins (Figure 2). In their absence, the green labeled nucleic acid was homogeneously distributed (Figure 2B), however, in their presence, the nucleic acid was massively recruited at the GUVs membrane interface (Figure 2C and D). As observed in the images, the recruitment was not uniform on the GUVs surface, and DNA-stained particles sizing 1-2µm diameter were either bound to or surrounded by membranes (Figure 2C, D,). In addition, when labelled LUVs and ssDNA were present, the nucleic acid and liposomes were both targeted to the GUVs interface mediated by the capsid proteins (Figure 2G and H; Figure 2F: control before protein addition).

**Figure 2.**
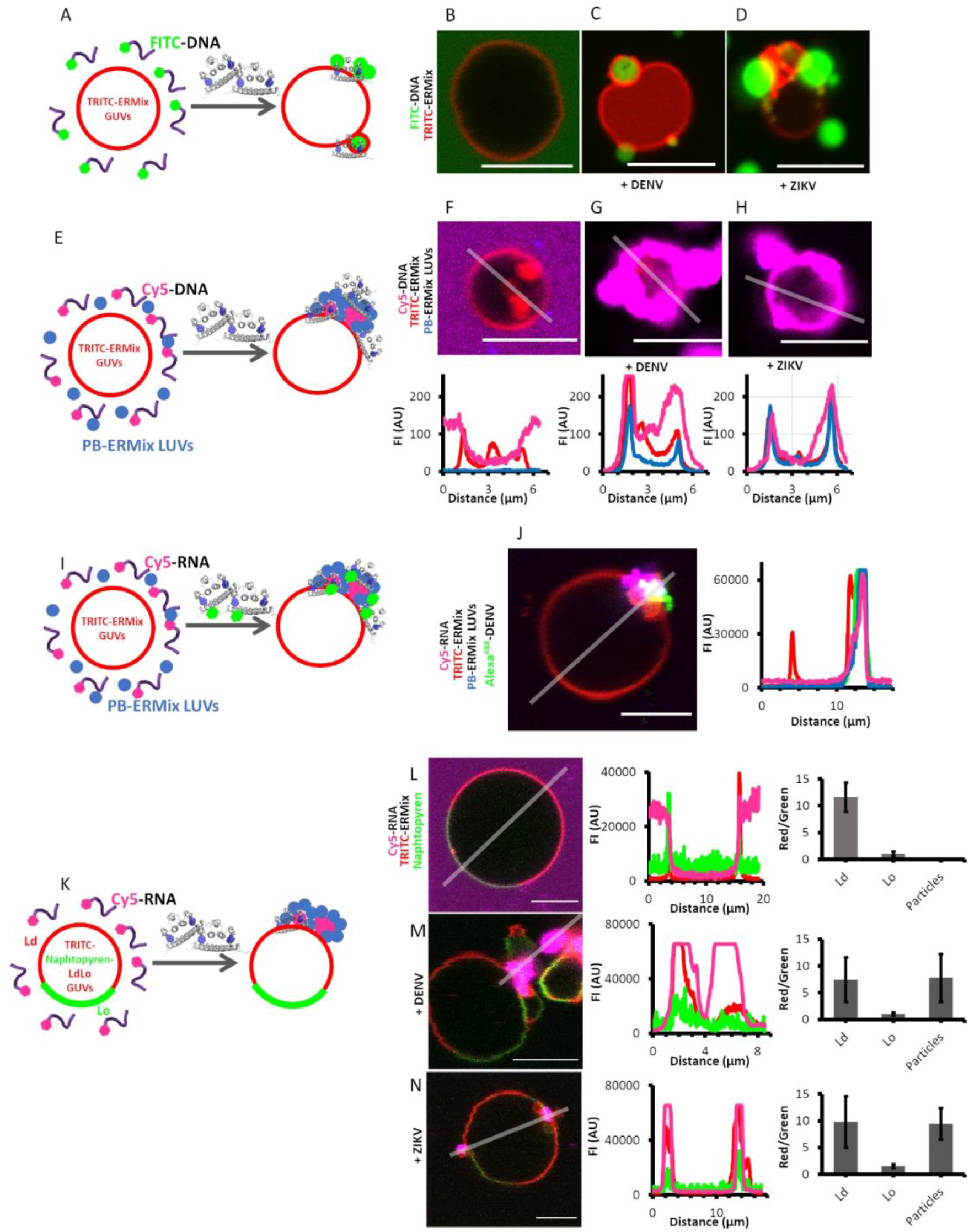
DENV and ZIKV capsid interact with ERmix GUVs and recruit to the lipid membrane LUVs and oligonucleotides. Schematic representation of FITC labelled ssDNA recruitment at the membrane of ERmix GUVs doped with TRICT-PE after capsid protein addition (A). Representative color merged confocal fluorescence images of TRITC-PE ERmix GUVs (red channel) incubated with FITC-DNA (green channel) in the absence (B) or in the presence of DENV (C) or ZIKV (D) capsids denote the particular recruitment of the labelled oligonucleotide by the capsids. Protein-triggered liposome and oligonucleotide recruitment to the GUVs membranes is cartooned in E. Representative color merged confocal fluorescence images of TRITC-PE ERmix GUVs (red channel) incubated with Cy5-DNA (magenta channel) and with PB labelled ERmix LUVs, in the absence (F, upper image) or in the presence of DENV (G, upper image) or ZIKV (H, upper image) capsids. The lower panels of F, G and H show the fluorescence intensity profile of the red, magenta and blue channels along the grey line. Alexa^488^-DENV also causes oligonucleotide and LUVs recruitment to the membrane of GUVs (I). Representative color merged confocal fluorescence images of TRITC-PE ERmix GUVs (J, red channel) incubated with Cy5-DNA (J, magenta channel) and with PB labelled ERmix LUVs (J, blue channel) after addition of Alexa^488^-DENV (J, green channel). The right panel in J, shows the fluorescence intensity profile of the red, magenta, blue and green channels along the grey line. DENV and ZIKV capsids recruit CY5-labelled RNA at Ld regions of the membrane of GUVs with Lo/Ld lipid phases (K). Representative color merged confocal fluorescence images of TRITC-PE (J, red channel, Ld marker) and naphthopyrene (J, green channel, Lo marker) labelled ERmix GUVs incubated with Cy5-DNA (J, magenta channel) before (L) and after addition of DENV (M) or ZIKV (N) capsid proteins. The middle panels in L, M and N show the fluorescence intensity profile of the red, magenta and green channels along the depicted grey line. Quantitative bar plots at the right panels of L, M and N show the red/green fluorescence intensity ratio for Ld and Lo regions and that at the membrane region where Cy5-RNA particles are observed. Note that some fluorescence signal profiles plotted reach their maximum values because image are acquired with exactly same settings as the control situation. Results are from at least two independent experiments (quantitative plots showing average and standard deviations) and images show representative observations. Scalebar in confocal images scales to 5µm.

Furthermore, to better characterize the system, the capsid proteins were labeled. Adding Alexa^488^-cysDENV to the co-suspension of ERmix GUVs, Cy5 fluorescently labelled RNA and small PB stained LUVs, the same phenomena was observed at the GUVs interface, confirming that the viral proteins trigger and are contained in these supramolecular aggregates (Figure 2I and J). These observations are particularly relevant because they indicate that DENV and ZIKV capsid proteins can spontaneously recruit nucleic acids and membranes onto the interface of a targeted membrane.

Finally, when GUVs display Lo/Ld domains, the RNA/DNA-containing particles are localized at regions corresponding to liquid disordered phases (red channel, Figure 2L, M and N), suggesting a site-specific membrane association when capsid proteins recruit RNA. The experiments highlight that the process involves a large amount of material clustered as a puncta or droplet shape and is always enhanced when nucleic acids are present. These aggregates resemble liquid condensates as reported for several similar situations when positively-charged (proteins) and negatively-charged (nucleic acids) polymers interact (13–15).

### DENV and ZIKV conform reversible liquid condensed droplets when interact with RNA

When nucleic acids interact with the viral capsid proteins, we regularly observed an increase of turbidity and spherical aggregates attached to the glass, (like those noticed to be associated to GUVs). Particles showing either DNA or RNA can be observed thorough the fluorescence microscope showing a broad size distribution (Figure 3A and B left and middle panels). DENV particles interacting with Cy5-DNA present an average diameter of 0.87 µm while, in the same conditions, ZIKV particles are 0.77 µm. Comparing their size and shape these type of condensates correlate to self-assembled nucleic acid-driven liquid droplets (3). Therefore, we further explored the phase-separation process and analyzed the reversibility and physicochemical properties of the new phase. For example, the aggregate assembly can be reverted by increasing the ionic strength of the environment. This is shown in the quantitative bar plots at the right panel of figure 3A and B, where at low NaCl concentration (<150 mM) the turbidity of the samples containing either DENV or ZIKV capsid proteins and non-labelled DNA (25mer) is high, indicating the presence of particles that can scatter light, while this effect is diminished above this concentration. Control experiments (Figure S1) with heparin, a well-known liquid droplet stabilizer (16), are in line with the salt-dependent reversibility of the phase separation induction by polyanionic molecules (i.e. RNA or DNA). Once formed, these self-assembled drops can recruit an exogenously added protein. This is the case when Alexa^488^-cysDENV is injected into a chamber containing DENV/Cy5-RNA stabilized particles (Figure 3C). Not only all particles became labelled by the fluorescent version of the capsid (Figure 3C, left and upper right panel) but also some newly formed droplets stick together or appear from an already existing seed (Figure 3C, lower right panel). This kind of dynamism is a key characteristic for such entities, proposed to be a reservoir of several biomacromolecules (3).

**Figure 3.**
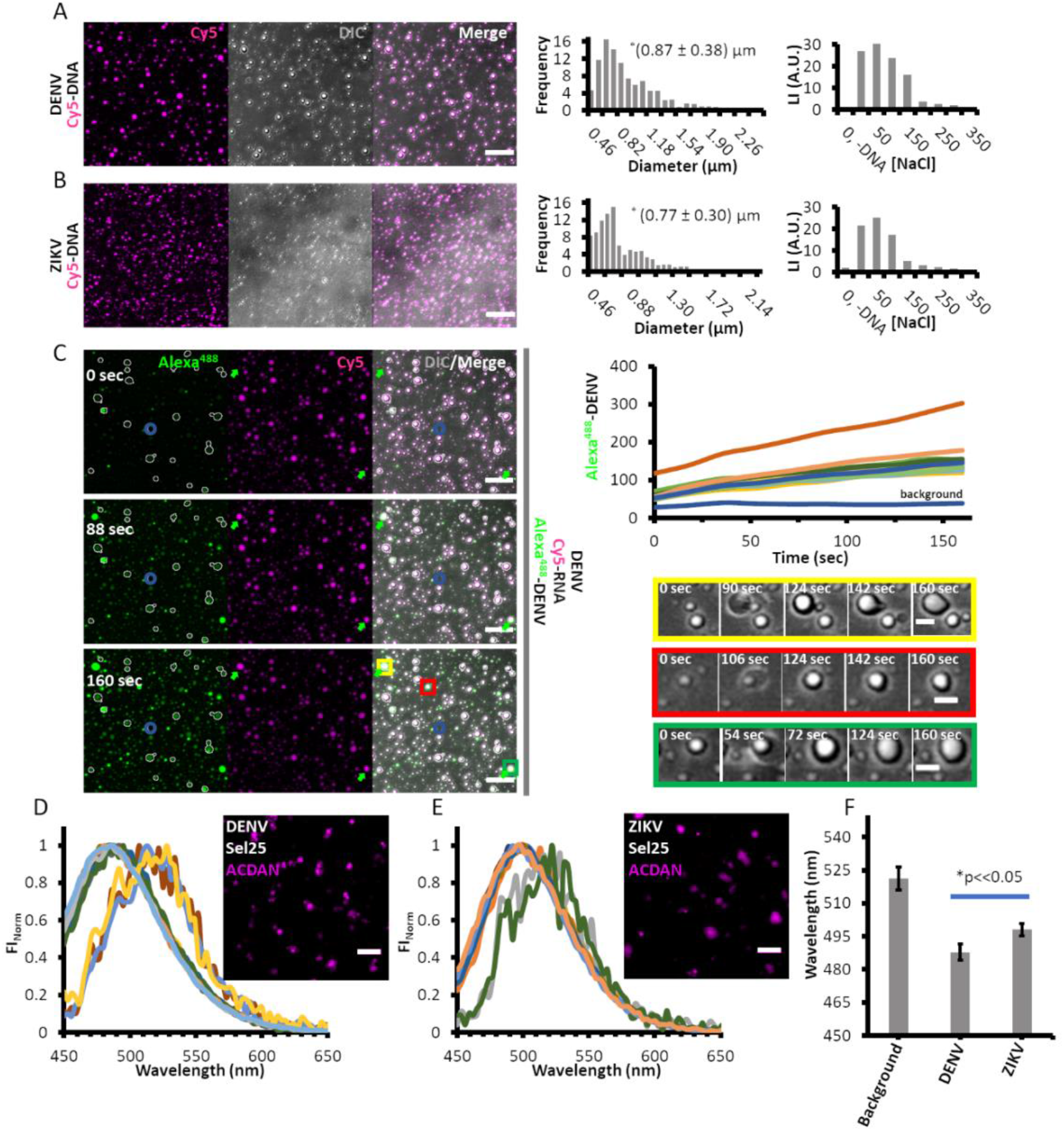
DENV and ZIKV capsid phase-separate into liquid droplets when interact with oligonucleotides. Representative confocal fluorescence images of Cy5-DNA (magenta) incubated with DENV (A, left panel) or ZIKV (B, left panel) from where the fluorescence oligonucleotide colocalizes with spherical particles observed by DIC (grey channel). From the images, the histogram of the particle size distribution frequency can be measured as represented in the middle plots of A and B. At the right panel of A and B, nephelometry measurements (see materials and methods methods) are reported showing scattered-light intensity versus NaCl concentration when DENV (A) or ZIKV (B) capsids phase-separate when interact with DNA depicting the reversibility of the phase-separation process. Incubation of DENV and Cy5-RNA (C, magenta) develops Cy5-positive droplets that can recruit exogenous injected Alexa^488^-DENV (C, green) in a time-dependent manner as shown in the upper right panel where the fluorescence intensity of Alexa^488^-DENV increases at the ROIs where Cy5 positive particles are present (C, upper right plot). From these representative experiments, several situations can be identified (yellow, red and green rectangles at the bottom right panel of C showing DIC images of the respective colored squares in the color-merge panel at the 160s time point) where new particles are generated and fuse with neighboring particles (yellow) or particles grow from a pre-existing seed (red and green). Water structure in the bulk or at the region of the droplets is measured with spectral confocal fluorescence microscopy (inserted images in D and E; see material and methods) taking advantage of the sensibility of the ACDAN dye to water dipolar relaxation. In this sense a spectral blue shift is observed for the emission spectrum of ACDAN colocalizing with DNA-stabilized DENV (D) or ZIKV (E) particles respect to the sample solution. Quantitative analysis of the spectral imaging (F) evidences a differential maximum emission wavelength of each condition (background: no particles and DENV or ZIKV droplets). Results are from at least two independent experiments (quantitative plots showing average and standard deviations) and images show representative observations. Scalebars in confocal images scale to 5 µm (A, C left panel; D and E) and 1 µm (C, bottom right panel). * in F represents a significative statistical difference between the compared values evaluated with a t-test of two-sample assuming equal variances (α = 0.05).

We also explored whether the extent of water dipolar relaxation is different in this new phase compared to that observed in bulk water. For this purpose, we used the polarity-sensitive dye 6-acetyl-2-dimethylaminonaphthalene (ACDAN), whose florescent properties largely depend on the dipolar relaxation of water in their vicinity (17, 18), a parameter connected to the rotational dynamic of water. Figures 3D and E show the change of the fluorescence emission spectrum of ACDAN at the interior of DNA-stabilized DENV and ZIKV particles, respectively, with respect to the solution, measured by spectral fluorescence microscopy. There is a noticeable shift of the ACDAN maximum emission intensity (Figures 3F) from 521 ± 5 nm (determined from ROIs where the dye is outside the particles) to 488 ± 3 nm (DENV) and 498 ± 2 (ZIKV) (determined from ROIs where the dye is at the particles), indicating an important change of the extent of water relaxation in the droplets. Of notice is the statistically significant difference of the values found for ZIKV particles compared to those from DENV, suggesting a differential capacity to affect water by each capsid conforming the droplets.

### Thermodynamics of the induced liquid-liquid phase transition when nucleic acid interact with DENV and ZIKV capsid proteins

To shed light into the thermodynamics of the liquid-liquid phase transition for the nucleic acid-capsid interaction, we took advantage of fluorescence anisotropy measurements of Cy5-labelled oligonucleotides. As previously reported, and increase in the turbidity of the sample can originate a decrease in anisotropy (19), an effect that can explain our results (homotransfer effects in the samples are discarded, see material and methods). Figure 4 summarizes the Cy5-DNA anisotropy change upon titration with DENV or ZIKV capsids. From these data, the fraction of adsorbed nucleic acid into the new phase is estimated. This allows, not only to calculate relevant thermodynamic parameters of the process (association constant, free energy), but also to understand its cooperativity by using the Hill equation (see materials and methods). Quantification from anisotropy data related to the concentration of DNA/RNA-protein complex formation ([LR]) upon protein addition (Figure 4 A, B, D and E) confirms that the nuclei acid-capsid association and the aggregation process is a cooperative event, since Hill coefficients (n) are higher than 1 (Figure 4C and F). We obtained apparent dissociation constants (Kd) in the order of 10^2^ nM, in agreement with those found, for example, for aptamers of proteins obtained by the SELEX methodology (20). From these data, the free energy of the transition process (ΔG_l-l_), both for DNA and RNA interacting with DENV and ZIKV capsids, can be estimated (Figure 4C and F). Our data shows an exergonic process taking place in both cases.

**Figure 4.**
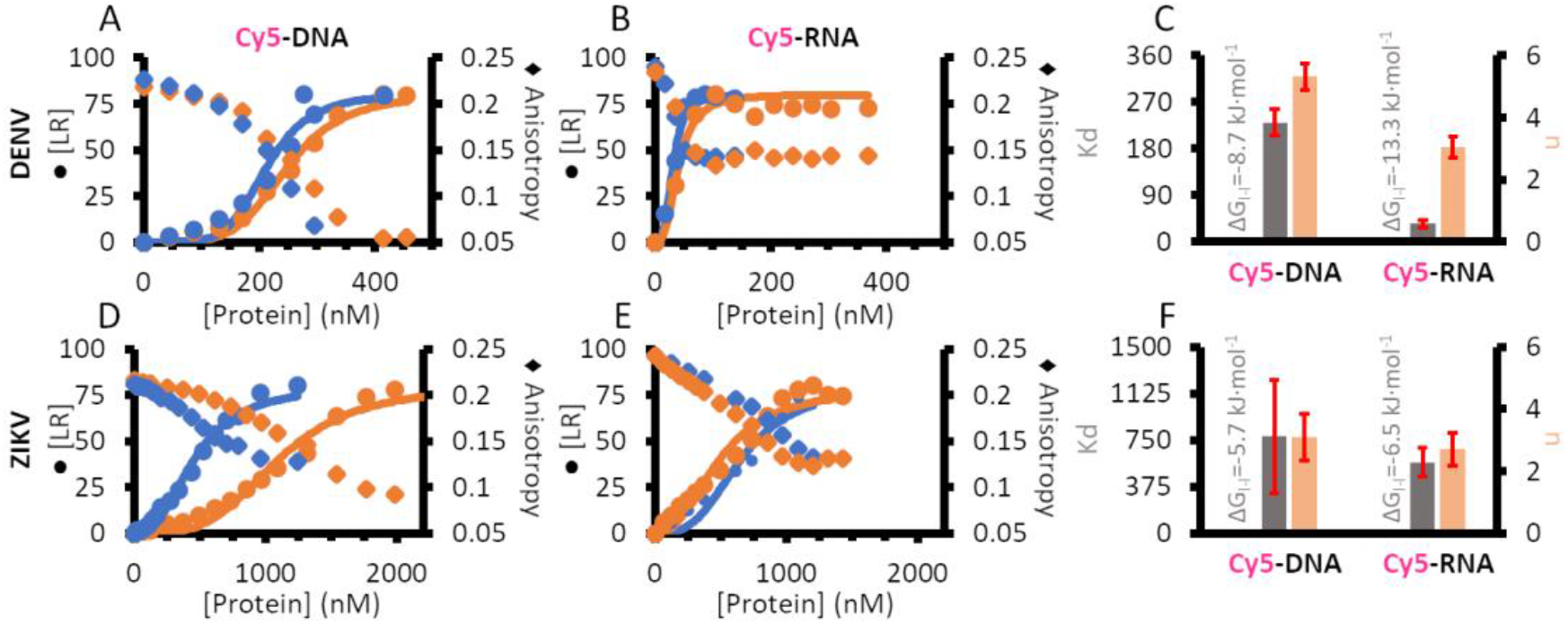
Thermodynamics of DENV and ZIKV interaction with oligonucleotides assessed by fluorescence anisotropy. Concentration of oligonucleotide-protein complex ([LR], circles) and anisotropy (diamond) changes of Cy5-DNA (A, D) and Cy5-RNA (B, E) in function of DENV (A, B) or ZIKV (D, E) concentration. Blue and Orange data represent different individual experiments and solid lines in A, B, D and E are the experimental data fitting to the Hill equation (see Materials and Methods section). From the fitting the dissociation constant, Kd (C, F; grey bars) and the Hill coefficient, n (C, F; light brown bars) are obtained for the interaction of Cy5-DNA (C and F, left bar set) and Cy5-RNA (C and F, right bar set) with DENV (C) or ZIKV (F) capsid proteins. Shown results are two independent experiments (quantitative plots showing average and standard deviations).

## Discussion

Maturation of Dengue and Zika viral particles at the membrane of the ER is a coordinated event where viral RNA is tightly packaged by capsids proteins and wrapped by the ER membrane before exiting the cell. So far, the NC approximation and ER association is proposed to be by protein-protein interaction with prM and E proteins at this organelle (5, 21). Recent data using ZIKV virus have shown that the transmembrane anchor peptide following the capsid protein may be partially retained on some molecules promoting oligomerization at the ER membrane, interaction with transmembrane regions of prM and E proteins, and consequent stabilization of particle assembly (22). Undoubtedly, the viral particle interior conforms a dense and condensed liquid phase as the result of the above-mentioned interactions. Comprehension on how this phase is stabilized and how it is specifically localized at the correct organelle is essential to characterize virus development and to approach crucial steps in where the viral morphogenesis can be blocked. In this sense, we studied how DENV and ZIKV capsids can interact with membrane model systems that mimic the ER lipid composition and the effect of the presence of DNA/RNA molecules and small liposomes along this association.

We find that both viral capsids can bind liposomes to promote membrane mixing, probably as a result of a membrane hemifusion events. Even more, this membrane recruitment is observed together with DNA/RNA binding onto the interface of giant liposomes. The associated capsids on the lipid bilayer dock both small LUVs and exogenously added nucleic acids, generating a massive local membrane concentration at the surface of giant liposomes. Of note is that the protein/nucleic acid/lipid condensations were localized at the liquid disordered domains of GUVs, showing a preferred lipid physical state for a potential viral particle formation. In addition, we describe how DENV and ZIKV capsids phase-separate when bind DNA or RNA. In this regard, we observed that the new phases display a droplet shape that can recruit extra-added material, such as the fluorescent version of DENV capsid or labelled nucleic acids. By analyzing the extent of water dipolar relaxation inside these droplets with ACDAN, we observed a slight spectral difference between DENV and ZIKV droplets. Specifically, the overall extent of water relaxation sensed by ACDAN in the case of DENV containing droplets is reduced respect to ZIKV case. These differences may be due to structural capsid dissimilarities (7–9) and, in consequence, compositional features of these two proteins, which differentially impact in their ability to structure water (23). In fact, ZIKV capsids seem to be more flexible than those of the DENV protein and this may impact on the water dipolar relaxation inside the capsid droplets. Furthermore, the thermodynamics of the liquid-liquid transitions reflects a spontaneous and a very cooperative process where a tight interaction of the oligonucleotides (in the order of 10-10^2^ nM) is observed.

Altogether, we provide compelling data on the physicochemical properties of complexes that include membranes, viral proteins and nucleic acids, proposing a new role for liquid droplet formation and membrane docking on the viral nucleocapsid assembly, as schematized in Figure 5. Considering the virus encapsidation process not as a result of mere molecular interactions but as an emerging new liquid phase might help to clarify other important aspects of the virus development. For example, the nucleocapsid structure also have a role in protecting the integrity of the viral genome, which is very important for the virus survival. To phase-separate the viral RNA inside the cell can contribute to exclude cellular components that would compromise its stability and avoiding damage. Actually, the fact that the water activity is different in this new phase, as we show here, could have as a consequence a possible reduction of water-dependent enzymatic degradation, for instance, by hydrolytic RNases (24, 25). Therefore, understanding the capsid capacity to bind membranes and to recruit genomic information into the lipid interface in a new liquid phase, a novel scenario of possible targets to impair viral particle maturation is open. In this sense, the development of new antiviral therapies can be focus on the disruption of the capsid-ER or capsid-RNA association and liquid condensation.

**Figure 5.**
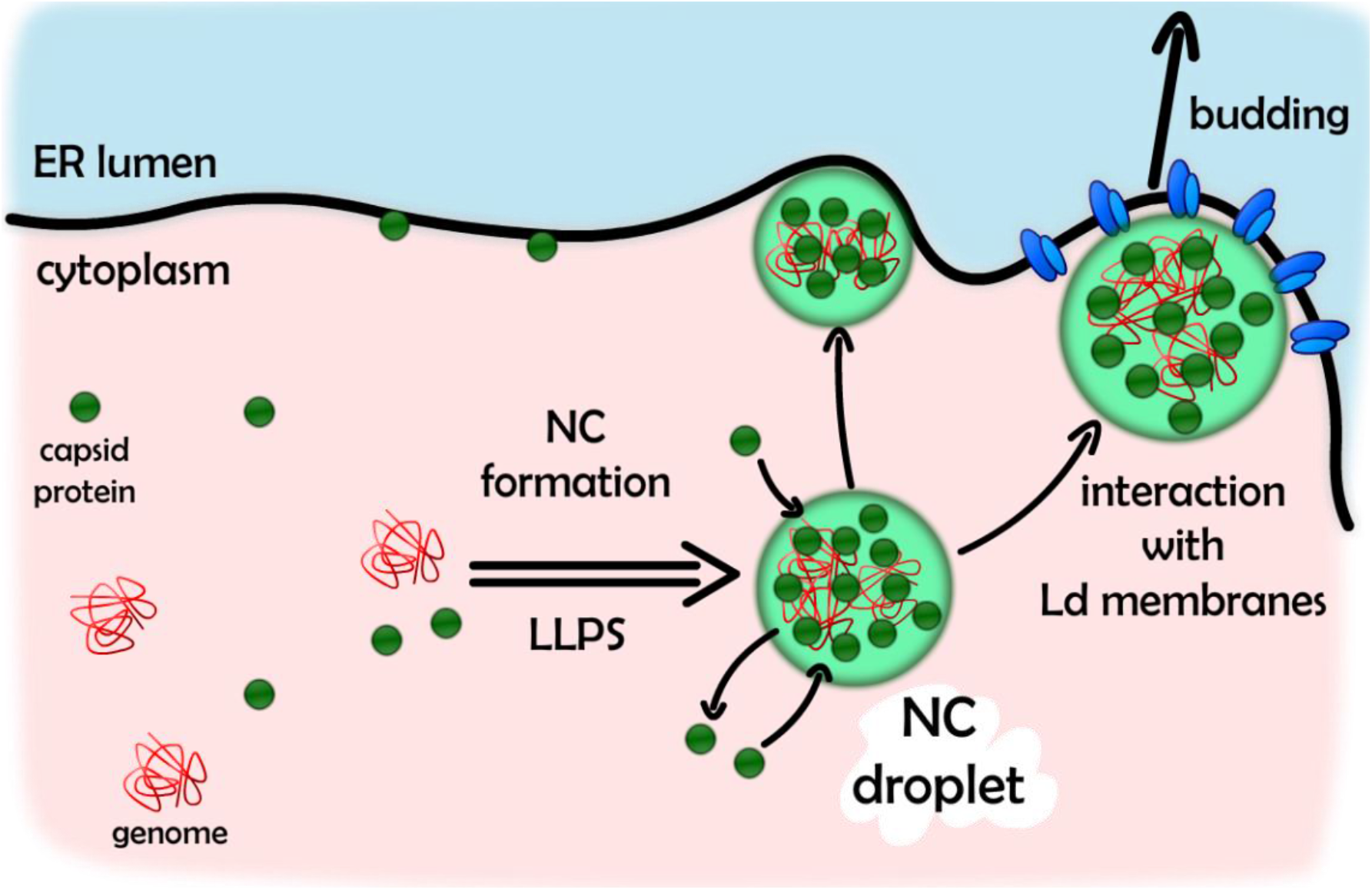
Schematic representation of the encapsidation indicating a process of liquid-liquid phase separation (LLPS). The interaction between capsid protein (green circles) and the viral genome (in red) results in a liquid-liquid phase transition to form the NC structure (light green droplet). These droplets associate with the ER membranes during the viral particle formation stabilizing a nucleocapsid-membrane interaction for final viral particle budding into the ER lumen.

## Experimental procedures

### Reagents

Oligonucleotides ssDNASel25 (5’GGACAGGAAUUAAUAGUAGCUGUCC3’) and FITC(5’)-ssDNASel25 oligonucleotides were obtained from Invitrogen (USA), Cy5-DNA (5’ Cy5-AAACTATTATTACTCATTTTCCGCCAGCAGTCAACTTCGATTTAATTCGTAAACAGATCT3’) was purchased from Macrogen (Korea) and Cy5-RNA (5’ Cy5-CAUCAUGCAGGACAGUCGGAUCGCAGUCAG 3’), Cy3 RNA (5’ CUGUCCUGCAUGAUG-Cy3 3’) and fRNA (5’ FAM-AGUUGAGUUGAGUUG 3’) were purchased from IDT (USA). For the experiments presented in this work, the oligonucleotides were diluted in DNase-free and RNase-free water.

All lipids used in the experiments (10 mg/mL) including Egg NBD-PE (L-α-Phosphatidylethanolamine-N-(7-nitro-2-1,3-benzoxadiazol-4-yl); 1mg/mL) were purchased from Avanti Polar Lipids, Inc. (Alabama, U.S.A.). TRITC-PE (N-(teramethylrhodamine-6-thiocarbamoyl)-1,2-dihexadecanoyl-sn-glycero-3-phosphoethanolamine, 1 mg/mL in 2:1 chloroform:methanol) is from Invitrogen (USA) and naphthopyrene from Sigma-Aldrich (St. Louis, MO). 6-acetyl-2-dimethylaminonaphthalene (ACDAN) was purchased from Santa Cruz Biotechnology Inc. (Dallas, TX). All stock solutions were prepared in chloroform:methanol 2:1 (v:v) at concentrations ranging 20 to 1 mg/mL and kept at -80 °C. All laboratory reagents (including buffers and salts) are from Sigma and Merk (USA) and used without further purification. Water was either milli-rho or milli-Q quality, obtained from an ion-exchange resin.

### Proteins

Constructs encoding Capsid proteins were derived from the cDNA of an infectious clone of DENV2 strain 16681 (GenBank accession number U87411) and the cDNA of a lab-adapted virus from Senegal, A1C1/V2 (GenBank accession number KX198134) from Zika virus (ZIKV). The coding sequence of DENV capsid protein was amplified from residues 1 to 100 by PCR and cloned into a pET15b vector between NcoI and BamHI to generate plasmid pET-DENVC. The coding sequence of ZIKV capsid protein was amplified from residues 1 to 104 by PCR and cloned into a pET15b vector between NcoI and BamHI to generate plasmid pET-ZIKVC.

For protein expression, competent Rosetta BL21 (DE3 pLys) E. coli bacteria were transformed with the above-mentioned expression plasmids. Transformed cells were grown at 37°C in LB medium with ampicillin (100 µg/mL) and chloramphenicol (25 µg/mL) to an optical density of 0.6 (600 nm wavelength), and then induced with IPTG at a final concentration of 0.5 mM. The induced cells were incubated at 20°C for 5 hours and the cells were then harvested by centrifugation at 4000rpm for 10 minutes. The cell pellet was resuspended in lysis buffer containing 50 mM NaH_2_PO_4_/Na_2_HPO_4_, 0.1 M NaCl, 20% glycerol, 1% Triton X-100, protease inhibitor cocktail (Sigma), 5 mM 2-mercaptoethanol. The suspension was sonicated, clarified by centrifugation, and then filtered with a 0.45 µm syringe filter. The cell extract was applied to a HiTrap Heparin HP column (GE Healthcare) equilibrated in 50 mM NaH_2_PO_4_/Na_2_HPO_4_ buffer with 2% glycerol and washed and eluted with the same equilibration buffer with increasing concentrations of NaCl. The fractions containing the capsid protein were pooled and concentrated using an Amicon Ultra-15 centrifugal filter (Merck Millipore), and the buffer was exchanged to a final storage buffer composed of 50 mM NaH_2_PO_4_/Na_2_HPO_4_. The protein was stored at -20°C.

A cysDENV (N93C) cysteine mutant of DENV capsid was generated by replacing the asparagine at the position 93 by cysteine. The mutation was incorporated in the reverse primer used to amplify the coding sequence of DENV capsid protein by PCR. The mutated protein was cloned, expressed and purified as previously described. This mutant allows us to label the capsid protein with a maleimide-derivative dye. In our case we used Alexa^488^-maleimide to label cysDENV and to be able to detect the protein directly thorough the fluorescence microscope. Protein labelling was carried out by adding 20 µL of 0.1 mg/mL Alexa^488^-maleimide in DMSO into a cysDENV aliquot, to a final dye/protein ratio of 5. The suspension was incubated up to 1 h and the non-coupled free dye was removed by exclusion chromatography using a PD10 exclusion column (GE Healthcare, Sweden) equilibrated with 50 mM NaH_2_PO_4_/Na_2_HPO_4_ with 1% glycerol.

### Liposomes

Liposomes were mainly composed of a lipid mixture (ERmix), resembling the ER lipid composition, following that reported by Keenan and Morré, 1970 (26). Specifically, the composition of the ERmix lipid used was (mole %): 4% brain sphingomyelin, 60% egg phosphatidylcholine, 3% liver phosphatidylserine, 9% liver phosphatidylinositol, 9% liver phosphatidylethanolamine and 15% cholesterol.

Giant unilamellar vesicles (GUVs) were obtained following the procedure described by Weinberger et al., 2013 (27). Briefly, 200 µL of a 5% (p/v) of PVA (polyvinyl alcohol; water solution) were spread onto a clean coverslip and dried for 1 h at 70 °C. Coverslips were cleaned using a 1% Hellmanex II (Sigma-Aldrich, USA) water solution for 3 hours. Once dried, 20 µL of the specific lipid mix in chloroform:methanol 2:1 at 0.5 mg/mL and doped with 0.5 to 2% fluorescent lipophilic dyes (i.e., TRITC-PE and naphthopyrene) were dispersed onto the PVA film with a Hamilton syringe in a dropwise manner. To eliminate the organic solvent, the sample was first evaporated with under a N_2_ stream followed by a treatment in a high vacuum chamber for 2 hours. GUVs were prepared by addition of 200 µL of a sucrose solution on the surface of the dried polymer cushion which contain the lipids. Osmolarities of the sucrose solution ranges from 290 to 310 mOsM as checked in a freezing point osmometer from Löser Messtechnik (Germany). After 45 min of the sucrose addition, GUVs are harvested by aspiration with a micropipette, kept in a plastic microtube and used the same day.

For the preparation of large unilamellar vesicles (LUVs) (28), the appropriate amount of lipids were mixed in conic glass tubes, slowly dried with a steam of nitrogen gas and left in vacuum for 2 – 3 hours. Dried lipids were supplemented with HKM buffer to a final total lipid concentration of 4 mM. This lipid suspension was frozen and thaw five times using N_2_ (l), and a thermal water bath set to 37 °C, respectively. After the last freeze-thaw cycle, the sample was extruded 20 times through a 100 nm pore size polycarbonate membrane from Whatman (Schleicher & Schuel) at 23 - 25ºC in Avestin extrusion device (Ottawa, Canada).

### Techniques

#### Fluorescence measurements

##### Interactions of DENV and ZIKV capsids with LUVs

All fluorescence measurements of this section were conducted on a Cary Eclipse Spectrofluorometer (USA) using a 3 mm optical path length cell and at 23-25 °C. The tryptophan (W) fluorescence emission spectra of the proteins, in the absence and in the presence of ERmix LUVs (1mM), were obtained from a 5 μM protein solution in 20 mM Hepes buffer with 120 mM potassium acetate and 1 mM MgCl_2_ (HKM buffer). W excitation was at 280 nm wavelength and the emission spectra observed at the 310-420 nm range.

To study the binding of the capsid proteins to LUV, FRET experiments between the protein W and the lipophilic probe NBD-PE were performed according to (12). Briefly 200 μM ERmix LUVs suspension, doped with 2% NBD-PE, in HKM buffer was titrated with different amounts of the proteins and the NBD fluorescence emission (acceptor) collected at 530 nm exciting W at 280 nm (donor fluorophore). The maximum protein to total lipid mole ratio reached is 0.05.

To evaluate a potential exchange of lipids among different liposomes, promoted by the interaction of the proteins with the membranes, we use another FRET couple. Briefly, 0.2 mM ERmix LUVs presenting 2% mole NBD-PE (donor fluorophore) and 3% mole TRICT-PE (acceptor fluorophore) were mixed with 0.8 mM of non-stained LUVs in HKM buffer. NBD fluorescence emission was registered at 530 nm with an excitation wavelength of 467 nm. If lipid from labelled and non-labelled liposomes are mixed upon protein addition, the donor-acceptor collisional probability decreases resulting in a net donor fluorescence increase (29). Experiments were performed by recording the donor signal in absence and presence of either 5 μM DENV or ZIKV capsid proteins (0.005 protein to total lipid mole ratio). Control experiments representing a total separation of the NBD-TRICT-PE FRET couple were performed by addition of triton X-100 (0.5% v/v final concentration).

##### Nucleic acid-protein interaction characterization by fluorescence anisotropy

Steady-state anisotropy measurements of Cy5-labeled DNA or Cy5-RNA were carried out on an ISS PC1 spectrofluorometer (ISS Inc., Champaign, IL). Excitation of Cy5 was performed with a 635 nm laser diode, and emission was collected through a 720 long pass filter. The excitation and emission slits used were 1 nm. At the maximal protein concentration used (up to 2.5 µM), no significant background due to scattered light or buffer (HKM buffer) fluorescence is detected. Every anisotropy point, from free oligonucleotide (80 nM) until the last protein addition, results from the average of 10 continuous measurements at 22 °C.

The gaseous equilibrium adsorption process between two phases and the molecular aggregation effect on the ligand binding were described by the seminal works of Irvin Langmuir and Archibald Hill, respectively (30, 31). We used the Hill adsorption isotherm to analyze the thermodynamics and the cooperativity of oligonucleotide-protein interaction (31). The fraction, *θ*, of ligand binding sites on the protein that are occupied by a ligand (L) is described as:

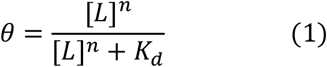

Where *n* is represented to be the binding site number but, experimentally, is a measure of the degree of interaction between these sites and [L] the concentration of free ligand (32). *θ* can be related to the concentration of ligand-receptor complex ([*LR*]) as 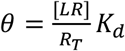, being *R*_*T*_ the total protein concentration. With this expression, a plot of [*LR*] vs. protein concentration is used to fit the model:

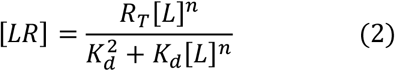

From our anisotropy data, we estimated [*LR*] as the total oligonucleotide *L*_*T*_ minus the free ligand fraction determined 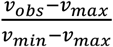L_T_, where *v*_*obs*_, *v*_*min*_ and *v*_*max*_ are the observed, minimum and maximum anisotropy values, respectively. Equation 2 was fitted to the experimental data by using OriginPro 8 (OriginLab Corporation, Northampton, MA, USA).

##### Fluorescence confocal microscopy and image analysis

Samples were studied with spectral Olympus FV1000 or FV1200 laser scanning confocal microscopes equipped with different lasers lines (FV1000: 405, 488, 561 and 633 nm; FV1200: 405, 473, 559 and 635 nm) and objectives (FV1000: 63X, 1.4NA; FV1200: 60X, 1.42NA) in a room with a temperature set at 23 °C. In general, sequential images at a fixed z position were acquired. Samples were observed either in an 8-well Lab Tek chamber slides (Promega, Madison, WI, USA) or a home-made Ludin chamber. Before observation, coverslips were passivated with casein 1% in HKM buffer for 20min and rinsed with HKM.

Experiments with GUVs were normally carried out by adding 1-2 μL of the stock of GUVs into a 0.6 mL plastic microtube then the incubated with 0.5 to 3 μM of proteins in KKM buffer. Depending on the experiment, a sequential addition of the other reagents (oligonucleotides and/or LUVs) was done. The working final volume was always 200 μL. The sample was added into the microscope observation chamber and GUVs were imaged after 15 min to ensure liposome sedimentation because of their sucrose content.

The time-sequence of the incorporation of Alexa^488^-DENV into pre-existing liquid droplets was conducted by incubating 3 μM DENV + 3 μM Sel25 DNA for 15 min (droplet formation) directly in the observation Ludin chamber, starting the time-lapse fluorescence confocal acquisition of 4 z-planes with a 0.5 μm z spacing and then injecting 8 μL of Alexa^488^-DENV 12 μM (reaching 0.5 μM final concentration). Every time point (4 sec) is the built up from the maximum intensity image projections. Once the time stack image was obtained, we used the particle analysis tool of the ImageJ software, thresholding the stack to individualize the 90% of the particles and to quantify the Alexa^488^ time-dependent signal increase at every single droplet.

Information of water dynamics at the oligonucleotide-protein stabilized droplets was obtained using the fluorescence probe ACDAN. As previously described (17) this probe responds to the extent of water relaxation occurring in its milieu. Specifically, as a first step we incubated either 3 μM DENV or 3 μM ZIKV with 3 μM Sel25 DNA for 15min directly in the observation chamber (200 μL). Then the fluorescent ACDAN dye was added from an ethanolic stock solution (1.4 mM) to a final concentration of 14 μM. The confocal spectral imaging experiments were performed by exciting at 405 nm and collecting the emission light between 450 and 732 nm with a resolution of 3 nm. For these experiments we used clean non-casein passivated coverslips in order to reduce possible interactions between ACDAN and the coverslip surface. Fluorescence emission spectra obtained from the spectral images at selected regions of interest were normalized to allow better comparison.

## Acknowledgments

This work was supported by grants from the National Agency for Research and Innovation of Argentina PICT 2015-2575 and PICT 2018-3204 to E.E.A and PICT-2019-02869 to AVG, and R01.AI095175 from NIAD to AVG. L.B.P.S. and GCN are supported by CONICET fellowships. E.E.A. L.A.B. and AVG are CONICET Investigators. All images were taken with microscopes from CEMINCO (Centro de Microscopia y Nanoscopia de Córdoba), Argentina. We are also grateful to all the members of the biophysical area at CIQUIBIC-DQB for helpful discussions.

## Author Contributions

E.E.A., L.A.B. and A.V.G. designed research; E.E.A., G.S.C.N, L.B.P.S. performed research; E.E.A., L.B.P.S., L.A.B., G.S.C.N, and A.V.G. analyzed data; and E.E.A., L.A.B., G.S.C.N, and A.V.G wrote the paper.

## Funding

This work was supported by grants from the National Agency for Research and Innovation of Argentina PICT 2015-2575 and PICT 2018-3204 to E.E.A and PICT-2019-02869 to AVG, and R01.AI095175 from NIAD to AVG.

The content is solely the responsibility of the authors and does not necessarily represent the official views of the National Institutes of Health.

## Conflict of interest

The authors declare no competing interest and no conflict of interest.

